# Cohabitation Shapes Collective Decision-Making in Medaka Fish

**DOI:** 10.1101/2022.06.14.494464

**Authors:** Ryohei Nakahata, Hideaki Takeuchi

## Abstract

Cohabitation within groups promotes behavioural synchrony and facilitates information transfer. Whether it shapes collective decision-making under predator threat is unknown. Here groups of six medaka (*Oryzias latipes*) cohabited for one month were used to test whether cohabitation promotes instantaneous collective decision-making in response to a looming stimulus (LS) mimicking a predator attack. Firstly, we analysed behavioural transitions before, during, and after LS in groups composed of six individuals. Individuals showing high-speed swimming in response to LS typically tended to shift to freezing-like behaviour afterwards, whereas non-responders were more likely to maintain normal swimming. Group-level analysis revealed a bimodal distribution in the number of individuals exhibiting freezing-like behaviour, with peaks at zero and six individuals, corresponding to ‘all non-freezing’ and ‘all-freezing’. Importantly, clustering analysis revealed that experimental groups showed consistent collective behavioural characteristics that could be categorised into three profiles: ‘all-freezing’, ‘all non-freezing’, and ‘partial freezing’. In non-cohabited groups assembled immediately before testing, the ‘all-freezing’ profile was absent, and the distribution in the number of individuals exhibiting freezing-like behaviour shifted to unimodal. In these groups, even at the individual level, responses more often showed a transition from high-speed swimming to normal swimming rather than freezing. The results indicate that cohabitation promotes synchronous freezing-like behaviour and consensus decisions under looming threat. Our study presents a behavioural assay for predator-evoked collective decision-making in a genetic model fish, providing a framework laying the groundwork for future efforts to link behavioural ethology with neuroscience.

## Introduction

Animals living in groups often exhibit synchronous behaviour in contexts such as migration, foraging, and predator avoidance. The emergence of coordinated behavioural patterns through interactions among individuals is termed collective behaviour, where group-level order and synchrony are thought to arise from local rules at the individual level, including attraction, alignment, and repulsion ^1^. Collective behaviour includes collective decision-making, in which a group selects a single option from among multiple alternatives, and this phenomenon is observed in various contexts such as movement^2,3^, foraging^4^, and predator evasion^5–7^.

In the context of predator avoidance, consensus formation during collective decision-making has been studied across various species. For instance, in sticklebacks, once the number of individuals escaping in a particular direction exceeds a threshold, the remaining group members tend to follow^6^ . Similarly, simulations involving humans have shown that when a critical number of escape responses are observed, the group tends to adopt avoidance behaviour ^5^. In elephants, the oldest female has been reported to be particularly sensitive to predator vocalisations and to influence the group’s decision to flee^7^. However, few studies have quantitatively investigated collective decision-making as an instantaneous group response under emergency conditions. In particular, how dynamic group-level responses to a rapidly approaching predator emerge remain poorly understood. While mechanisms underlying rapid individual decision-making in response to visual looming stimuli have been demonstrated^8^, systematic analyses of dynamic group-level responses remain scarce.

Cohabitation has also been shown to affect individual recognition and interaction patterns, thereby influencing behaviour and information transmission^9^. At the dyadic level, cohabitation enhances responsiveness to predators in predatory mites and reduces encounter frequency^10^. In sticklebacks, cohabitation reduced leadership tendencies in bold individuals, leading to more balanced coordination^11^. In cichlids, cohabitation promotes exploratory behaviour and reduces fear responses to novel stimuli^12^, while in zebrafish, the transmission of social fear is enhanced among cohabiting individuals^13^. At the group level, wild female guppies tend to associate with familiar cohabitants^14^, avoidance frequency in response to predator odour increases in head minnows^15^, and the latency to initiate avoidance of a predator model is reduced in brown trout^16^. In tropical damselfish, both responsiveness to fear stimuli and inter-individual information transmission are enhanced through cohabitation^17^. Overall, the literature indicates that cohabitation enhances alignment coordination and information transmission; however, how these factors influence collective decision-making remains unclear.

Furthermore, most empirical research in this field has relied on observations in natural environments or on wild individuals^6,15^, yet relatively few studies have been conducted in controlled experimental settings. Although collective decision-making regarding movement direction has been demonstrated in zebrafish^18^, reports of such decisions in response to predators are lacking. Moreover, integrative frameworks that link decision-making mechanisms at both the individual and group levels with molecular and neural analysis, particularly in genetically tractable model organisms, are still lacking.

To address these gaps, we focused on medaka *Oryzias latipes*, a well-established model organism in molecular genetics. Medaka are known to exhibit coordinated behaviour with conspecifics^19^ and to improve foraging efficiency through visual social learning^20^. Preliminary observations revealed that small groups of medaka responded synchronously to a human approach, either by showing ‘freezing-like behaviour after escape’ or by maintaining continuous movement without escape. Motivated by these findings, we aimed to develop a behavioural assay capable of quantitatively assessing instantaneous collective decision-making using a looming stimulus (LS) mimicking the sudden approach of a predator. LS has been used to elicit individual avoidance responses in mice^21^, zebrafish^22^, and fruit flies^23^. However, most previous studies have focused on individual-level decision-making^8^.Even in group contexts, the focus has remained on how individual responses are influenced by conspecifics ^23,24^, with little attention paid to whether the group as a whole converges on a single collective choice.

In this study, we optimised LS parameters and established a quantitative behavioural system capable of replicating the distinct response patterns during preliminary observations. We then tested whether medaka groups exhibit instantaneous collective decision-making in response to LS and examined the effects of cohabitation (joint rearing) on the decision-making patterns. Our findings establish an experimental model for examining collective decisions under acute threat. They also lay the foundation for elucidating how cohabitation modulates synchrony and group-level decision-making processes in a genetically accessible model organism.

## Methods

### Ethics statement

All the methods in this study were carried out in accordance with relevant guidelines and regulations. The work in this paper was conducted using protocols specifically approved by the Animal Care and Use Committee of Tohoku University (permit number: 2022LsA-003). All efforts were made to minimise suffering following the NIH Guide for the Care and Use of Laboratory Animals Fish and breeding conditions. The study was carried out in compliance with the ARRIVE guidelines (https://arriveguidelines.org/arrive-guidelines).

### Animals

Medaka (*Oryzias latipes*, fading strain) were obtained from Dr Tetsuro Takeuchi (Figure 1a)^25^. All individuals were hatched and bred in our laboratory. Medaka fish were maintained in groups in plastic aquariums (22.6 cm× 14.6 cm× 14.5 cm, Sanko) or custom-made acrylic aquariums(22 cm × 14.5 cm × 14.5 cm)under controlled temperature (26 ± 1°C) and light (14 h: 10 h light: dark) conditions. Every day, fish were fed brine shrimp between 12:00 and 13:00 and solid bait Otohime β-2 (Marubeni Nissin Feed, Tokyo, Japan) at least twice around 10:00 and 17:00 on weekdays. This study used individuals aged 2–9 months post hatch.

**Figure 1.**
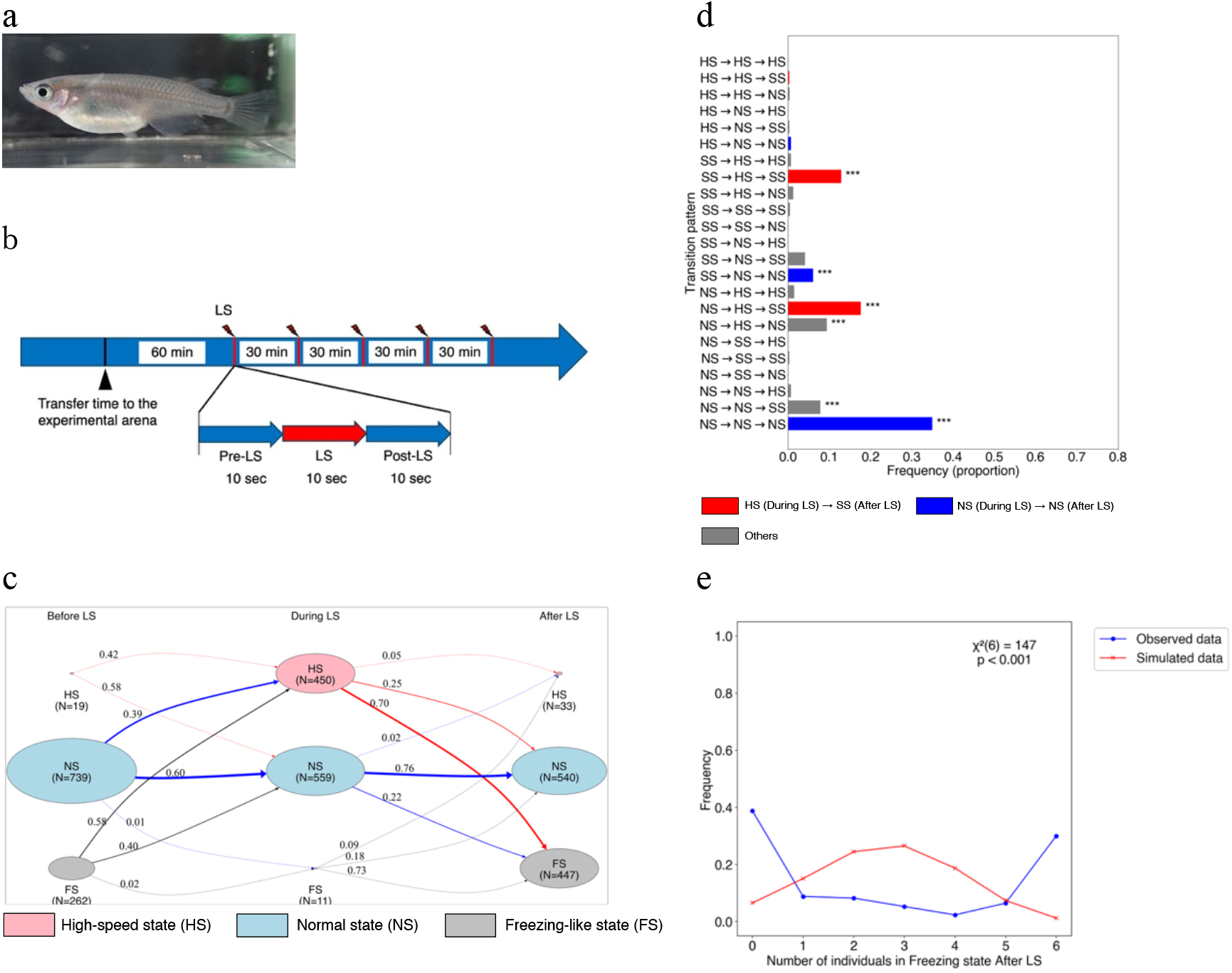
Experimental design, state transition analysis, and statistical evaluation of freezing-like behaviour in medaka after looming stimulus. a) Medaka fish (Fading strain) b) Groups of medaka were transferred, with their tanks, from the circulating water system to the apparatus after feeding. One hour later, the looming stimulus (LS) was presented five times at 30-minute intervals. The experiment was conducted over two consecutive days, giving ten LS. For analysis, three 10-second periods (before, during, and after each LS) were used, totalling 30 seconds per trial. c) A state transition diagram visualises individual-level states (FS: freezing-like state, NS: normal swimming state, and HS: high-speed swimming state) across three intervals: before, during, and after LS. Node size represents the proportion of individuals, and numbers within nodes indicate counts. Node colours are red for HS, light blue for NS, and grey for FS. Numbers on edges indicate the transition probabilities, and edge thickness corresponds to the number of individuals. Edge colours indicate the originating state: red for transitions from HS, blue from NS, and grey from FS. d) A bar plot showing the frequencies of transition patterns across the three intervals. Each pattern is categorised according to the sequence of states. Red bars indicate transitions to HS during LS followed by FS after LS. Blue bars indicate individuals that remained NS during and after LS. Grey bars represent all other patterns. To assess statistical significance, a binomial test with false discovery rate (FDR) correction was applied. Patterns with q < 0.001 are marked with ^***^. e) The X-axis shows the number of individuals in FS after LS, and the Y-axis shows its frequency. The blue and red lines represent the observed data (17 groups) and the simulated data (17 groups × 1000 trials, seed = 1, …, 1000), respectively. A chi-square test result is shown in the graph (χ^2^ (6) = 147, p < 0.001).

### Cohabited conditions

Medaka (*Oryzias latipes*) aged either one month or nine months were randomly selected to form groups. Ten groups of six individuals each were formed from one-month-old fish (N = 60), and seven groups were formed from nine-month-old fish (N = 42). These groups were formed without regard to sex, because the sex ratio of one-month-old fish could not be reliably determined. In contrast, the nine-month-old groups were adjusted to achieve a 1:1 sex ratio. Each group of six individuals was maintained separately for a month. In total, 17 groups of sexually mature individuals aged either two months or ten months that had undergone cohabitation, were used for experiments.

### Looming stimulus (LS)

The looming stimulus (LS) was a visual stimulus mimicking an approaching predator, created using the “Zoom” animation function in Microsoft PowerPoint (Figure S1). The stimulus expanded to a width of 14.5 cm over 5.5 s, gradually darkened over 10 s, and remained black for 3 min (Figure S1). It was presented on an LCD monitor (EXLDH271DB, I-ODATA) mounted on the side of the aquarium (Figure S2). Both plastic and acrylic aquaria, also used as breeding tanks, were employed. Approximately 1–2 h after daytime feeding (12:00–13:00), the medaka groups were transferred to the behavioural testing apparatus along with their breeding tanks, and the water level was adjusted to 6 cm. The LS was presented five times at 30-minute intervals for two consecutive days, beginning 1 h after transferring the aquarium from the rearing system to the testing apparatus (Figure 1b).

### Behavioural recording and tracking

Behaviour was recorded from above using an action camera (M80 Air, Apexcam, or HERO8, GoPro) at a resolution and frame rate of 4K (30 fps) or 2.7K (50 fps). Recordings lasted 5 minutes, beginning 2 minutes before the LS and ending 3 minutes after. For analysis, a 30-s segment was extracted for each trial: 10 s before, during, and after LS (Figure 1b). Video files were extracted using QuickTime Player, converted to JPEG format using FFMPEG (v4.4.1), and subsequently converted to MP4 format at 5 fps. Tracking was performed using UMATracker^26^, and coordinate data were obtained with the UMATracker-Tracking tool, applying either the Pochi-Pochi (manual positioning) or Group Tracker GMM algorithm. Tracking errors, such as identity swaps, were manually corrected using UMATracker-TrackingCorrector.

To convert pixel values to centimetres, the number of pixels along the centre of the long side of the aquarium was measured in ImageJ, which was based on the actual inner length (20.5 or 20.0 cm). Velocity (cm s□^1^) was calculated from coordinate data (5 fps). A velocity matrix (6 individuals × 10 trials × 17 groups; 1020 × 150 frames) was compiled, and a moving average was applied (window size = 5) using pandas v1.4.0 to smooth short-term fluctuations.

### Behavioural experiment under non-cohabited conditions

Medaka aged 4 or 9 months, reared in a recirculating aquaculture system, were randomly selected. From the 4-month-old fish (N = 36), six groups were formed, and from the 9-month-old fish (N = 36), another six groups, each consisting of six individuals, yielded a total of 12 groups. The sex ratio in each group was adjusted to 1:1. Group formation took place in the morning, and experiments were conducted 1 to 2 h after the afternoon feeding. Subsequent procedures were conducted in accordance with those described in the cohabitation experiments.

### Statistical analysis

#### (1) Characterisation of state transition patterns at the individual level

To calculate the state transition probabilities for behavioural transitions before, during, and after LS, we constructed state transition matrices by counting the number of transitions between states and normalising each row, following established methods using Markov chain analysis^27,28^. To visualise the transition dynamics, we created state transition diagrams using python-graphviz v0.20.3, where each state (NS, FS, and HS) was represented by a node, with edges indicating transition probabilities.

Furthermore, we used a binomial test to compare whether there were significantly more specific state transition patterns in the series of flows from before LS to after LS. In the binomial test, we set the null hypothesis that ‘the 27 behaviour patterns occur with equal probability (1/27)’ and performed a one-sided test. To control for type I errors due to multiple comparisons, we applied FDR correction to the binomial test results.

#### (2) Analysis of group-level freezing-like states after LS

Based on the analysis of individual-level behavioural patterns, we next examined group-level freezing-like states after LS. For each trial, the number of individuals in the freezing-like state after LS was counted. To test whether synchronous freezing-like states occurred, virtual datasets were generated by randomly shuffling the states of each trial among groups (17 groups × 1,000 trials; seeds = 1, 2, …, 1,000). The proportions of individuals in each state across all trials were compared between the virtual datasets and the observational data using a chi-square test (scipy v1.10.1). The null hypothesis was defined as: “The presence or absence of the freezing-like state for each individual is independent, and synchronous freezing-like states for the entire group occur at random.”

#### (3) Classification of group response profiles

To classify these characteristics, we performed a principal component analysis (PCA) on 27 individual-level behavioural patterns from before to after the LS intervention. We then calculated the cumulative contribution rate and reduced the number of dimensions to the minimum required to explain >95% of the variance. To visualise the cluster structure, we further projected the PCA-reduced data using UMAP^29^ (umap-learn v0.5.7). Classification was performed using spectral clustering (scikit-learn v1.2.2), and the optimal number of clusters was determined based on the silhouette coefficient. This coefficient approaches 1 when intra-cluster cohesion and inter-cluster separation are high; therefore, the number of clusters yielding the highest silhouette coefficient was selected.

#### (4) Comparison of state transition patterns at the individual level between clusters

Differences in the frequency of state transition patterns between clusters were evaluated using binomial tests with FDR correction, as described above.

#### (5) Analysis of group-level freezing-like states

To examine differences in group-level freezing-like states across clusters and between cohabited and non-cohabited groups after LS, we applied a generalized linear mixed model (GLMM)^30^. The dependent variable was the number of individuals exhibiting the freezing-like state (0–6) within each group. Cluster identity and the presence or absence of cohabitation were included as fixed effects. Experimental group identity, trial number, and group identity were incorporated as random effects to account for repeated measurements and inter-group variability. The model assumed a binomial distribution with a logit link function, which is appropriate for categorical or count data with hierarchical structure. Analyses were performed in Python v3.8.12. We used pyper v1.1.2 to call R v4.1.2, and the lme4 and multcomp packages for model fitting and post hoc tests. Tukey’s method was applied for multiple comparison correction.

## Results

### Definition of individual-level behavioural states and classification of state transition patterns

To capture how individual fish responded to LS (looming stimulus) in terms of state transitions, we expressed behavioural responses as transition patterns across three intervals (before, during, and after LS). In total, 17 groups of six individuals each (N = 102) were tested. For each group, fish were transferred to the experimental arena and habituated for one hour, after which the looming stimulus was presented five times at 30-minute intervals over two consecutive days (Figure 1b). This protocol yielded a total of 6 individuals × 10 trials × 17 groups of individual-level datasets for subsequent analyses. For this purpose, we first defined behavioural types for each interval based on velocity data. Using histograms and kernel density estimation (KDE) curves of velocity, we categorised behaviour into three types: ‘freezing-like behaviour’, ‘normal swimming’, and ‘high-speed swimming’. The rationale for these definitions is as follows. Some individuals exhibited freezing-like behaviour after LS. The velocity histogram for the post-LS interval showed a bimodal distribution with a trough at approximately 0.2 cm/s (Figure S3a). Therefore, frames with speeds below 0.2 cm/s were defined as ‘freezing-like behaviour’. During LS, escape-like responses characterised by high-speed swimming were observed. Such behaviours were rarely seen before the LS onset. Comparison of KDE curves for the pre-LS and LS intervals revealed minimal overlap above 6 cm/s (Figure S3b). Thus, frames with speeds of 6 cm/s or higher were defined as ‘high-speed swimming’. Frames with velocities between 0.2 cm/s and 6 cm/s were categorised as ‘normal swimming’.

Based on these speed-based classifications, we then defined the behavioural state for each 10-second interval (before LS, during LS, and after LS). An interval was categorised as a ‘freezing-like state’ if freezing-like behaviour persisted for ≥ 8 seconds, and as a ‘normal state’ if freezing-like behaviour lasted for < 2 seconds. During LS, escape behaviour occurred rapidly. Therefore, if high-speed swimming (≥ 6 cm/s) was sustained for 0.2 seconds (equivalent to one frame at 5 fps), we defined this as a ‘high-speed state’. This threshold reflects the minimum temporal resolution required to identify continuous motion. Using these thresholds, each of the three temporal intervals was classified into one of the three behavioural states: ‘freezing-like state’, ‘normal state’, or ‘high-speed state’ (Figure S4).

### Detection of individual-level state transition characteristics

To examine how the behavioural states of individuals transitioned from before LS to during LS, and from during LS to after LS, we calculated state transition probabilities using a Markov chain and visualised them as a state transition diagram (Figure 1c). Between the pre-LS and LS intervals, 39% of individuals transitioned from the normal state to the high-speed state, whereas 60% remained in the normal state. Among individuals that entered the high-speed state during LS, 70% transitioned to the freezing-like state after LS. In contrast, individuals that remained in the normal state during LS had a 76% probability of continuing in that state after LS. These findings suggest that individuals showing escape-like behaviour during LS tended to transition into a freezing-like state after LS, whereas those unresponsive to LS generally maintained normal swimming.

To statistically evaluate trends in state transition patterns, we extracted behavioural state sequences across the three intervals (before, during, and after LS). Each sequence was expressed as a combination of the three defined behavioural states: high-speed swimming (HS), normal swimming (NS), and freezing-like states (FS), resulting in 27 possible transition patterns. We quantified the frequency of each pattern across trials (Figure 1d).

The most frequent transition pattern was the maintenance of the normal state throughout the three intervals (NS→NS→NS; blue; q < 0.001). The second most frequent pattern was NS→HS→FS (red; p < 0.001), and the third was FS→HS→FS (red; q < 0.001), both involving a transition to high-speed swimming during LS followed by freezing-like behaviour: NS→HS→FS (red; p < 0.001) and FS→HS→FS (red; q < 0.001). Additional significantly overrepresented patterns were NS→HS→NS (grey; q < 0.001), NS→HS→FS (grey; q < 0.001), and FS→NS→NS (blue; q <0.001), all of which exceeded the expected frequency under a uniform distribution (1/27).

Among these six prominent transition patterns, NS→NS→NS and FS→NS→NS represent non-reactive behaviours where individuals maintained or returned to the normal state during and after LS (blue). In contrast, NS→HS→FS and FS→HS→FS represent reactive responses, characterised by high-speed swimming during LS followed by freezing-like behaviour (red). These two reactive patterns accounted for 41% and 30% of all observations, respectively, and thus constitute the typical individual-level responses to LS stimulation. Notably, escape without subsequent freezing (NS→HS→NS; approx. 9%) and no initial response followed by freezing after LS (NS→NS→FS; approx. 8%) were also observed at appreciable frequencies.

### Population polarisation into synchronous freezing-like and non-freezing-like states after LS

To investigate whether medaka groups exhibited synchronous responses (either freezing-like or non-freezing-like behaviour) after LS, we counted the number of individuals exhibiting freezing-like behaviour in each trial. The distribution was bimodal, with peaks at 0 and 6 individuals (Figure 1e, blue line), suggesting that entire groups tended to respond uniformly.

To determine whether this distribution could be explained by chance, we generated a virtual dataset by randomly shuffling the individual-level freezing-like and non-freezing-like classifications within groups of the same size (Figure 1e, red line). This reconstructed the expected distribution under the assumption that individuals responded independently of one another. A chi-square test comparing the observed and expected distributions revealed a significant difference (χ^2^ (6) = 147, p < 0.001). This result suggests that the strong bias towards either all freezing or no freezing within groups after LS is unlikely to have occurred by chance alone. Instead, it indicates that individuals within a group reacted in a synchronous manner through social interaction.

### Group response profiles to LS classified into three types

During the behavioural experiments, we observed groups in which all individuals synchronous and exhibited freezing-like behaviour, as well as groups in which all individuals remained unresponsive and continued swimming. Moreover, the same groups tended to display similar response tendencies across repeated trials. Based on these preliminary observations, we hypothesised that groups exhibit consistent and characteristic behavioural tendencies, which we define as group response profiles. To evaluate this hypothesis, we classified groups according to their individual-level behavioural patterns. Specifically, behavioural data from 10 trials per group were aggregated, dimensionality reduction was performed using principal component analysis (PCA) followed by UMAP, and spectral clustering was applied to classify the groups.

PCA indicated that 16 dimensions were required to exceed a cumulative variance contribution of 95%, and this was adopted as the optimal dimensionality (Figure S5). The reduced data were then projected into two dimensions using UMAP, revealing a clear distinct group-level structures (Figure 2g). Spectral clustering, guided by the silhouette coefficient identified three as the optimal number of clusters (Figure S6). Accordingly, groups were classified into three clusters (Figure S7).

**Figure 2.**
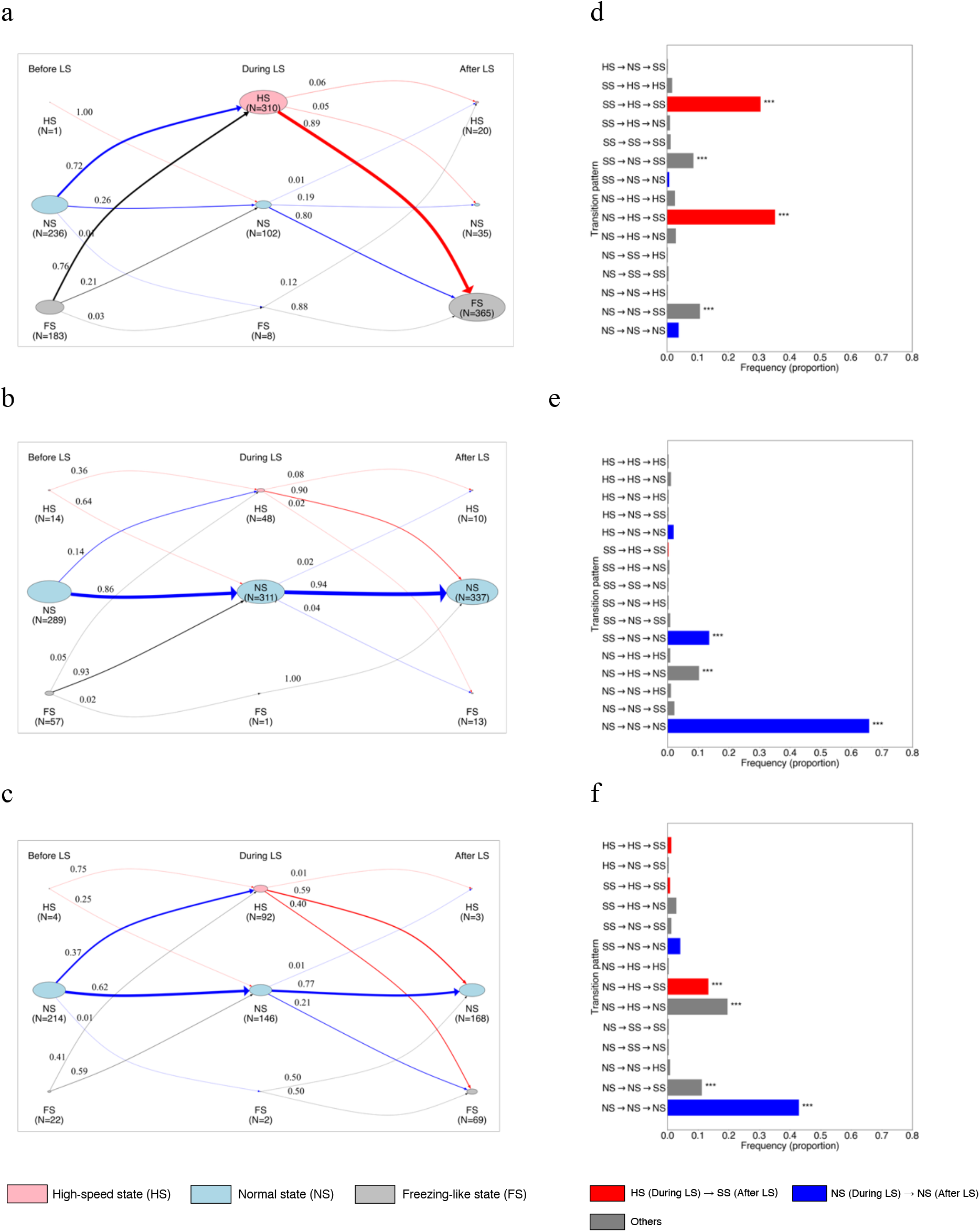
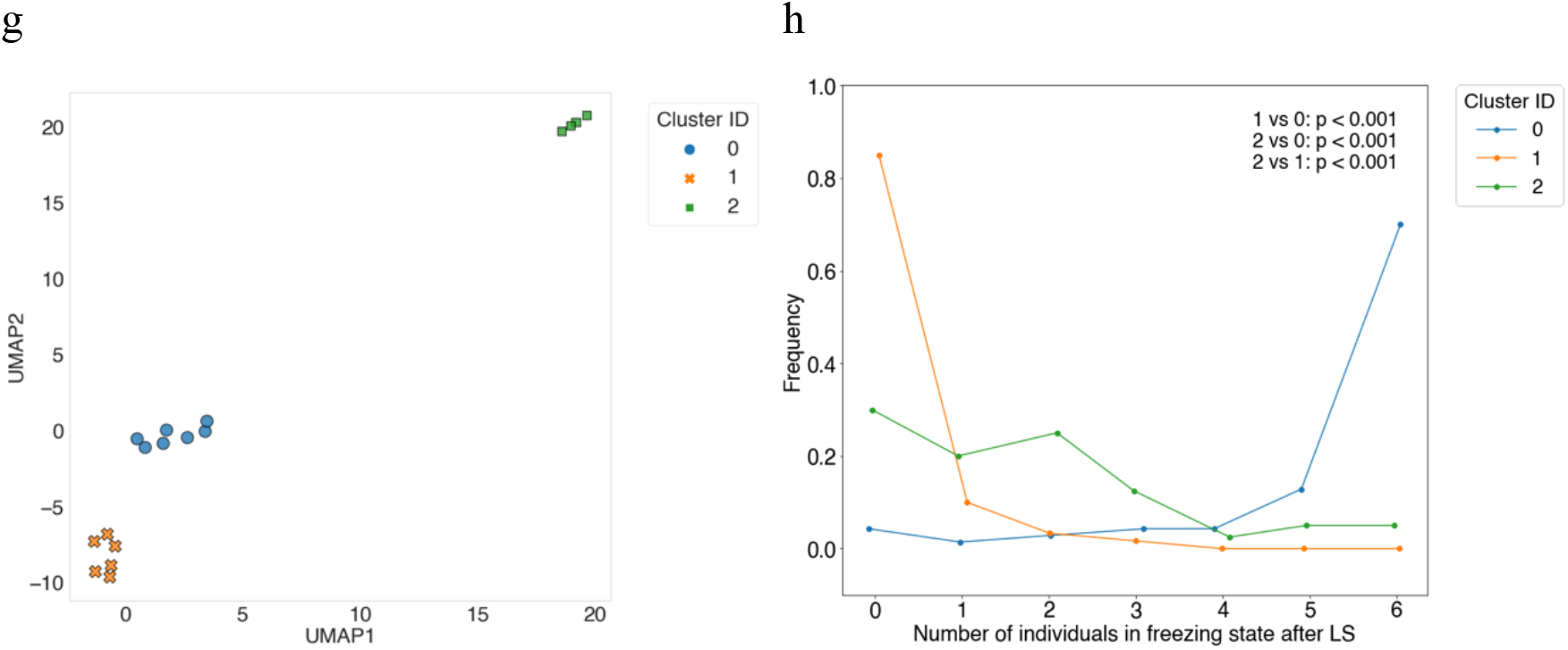
State transition diagrams and frequencies of behavioural state transition patterns for each cluster. a–c) State transition diagrams for each cluster were generated to represent individual-level behavioural states (FS: freezing-like state, NS: normal swimming state, and HS: high-speed swimming state) before, during, and after LS. All the visual elements are consistent with those in Figure 1c. a) State transition diagram for Cluster 0. b) State transition diagram for Cluster 1. c) State transition diagram for Cluster 2. d–f) Bar graphs showing the frequency of occurrence for each behavioural state transition pattern in each cluster. Details of colour coding and statistical tests are as described in Figure 1d. d) Distribution of transition pattern frequencies in Cluster 0. e) Distribution of transition pattern frequencies in Cluster 1. f) Distribution of transition pattern frequencies in Cluster 2. g) The X-axis represents the first UMAP component and the Y-axis the second. Each point shows the group centroid, obtained by reducing the original 23 dimensions to 16 dimensions using PCA and further to two dimensions using UMAP. Colours indicate classification results based on spectral clustering. h) The X-axis represents the number of individuals in the freezing-like state after LS, and the Y-axis represents the frequency of these counts across all trials. The lines correspond to the IDs of the three clusters. Tukey’s post hoc test based on a GLMM was performed, and the results of the cluster comparisons are shown within the graph.

To analyse state transitions of individuals across the pre-, during-, and post-LS intervals in each cluster, we calculated state transition probabilities using a Markov chain and visualised them as state transition diagrams (Figure 3a-c).

**Figure 3.**
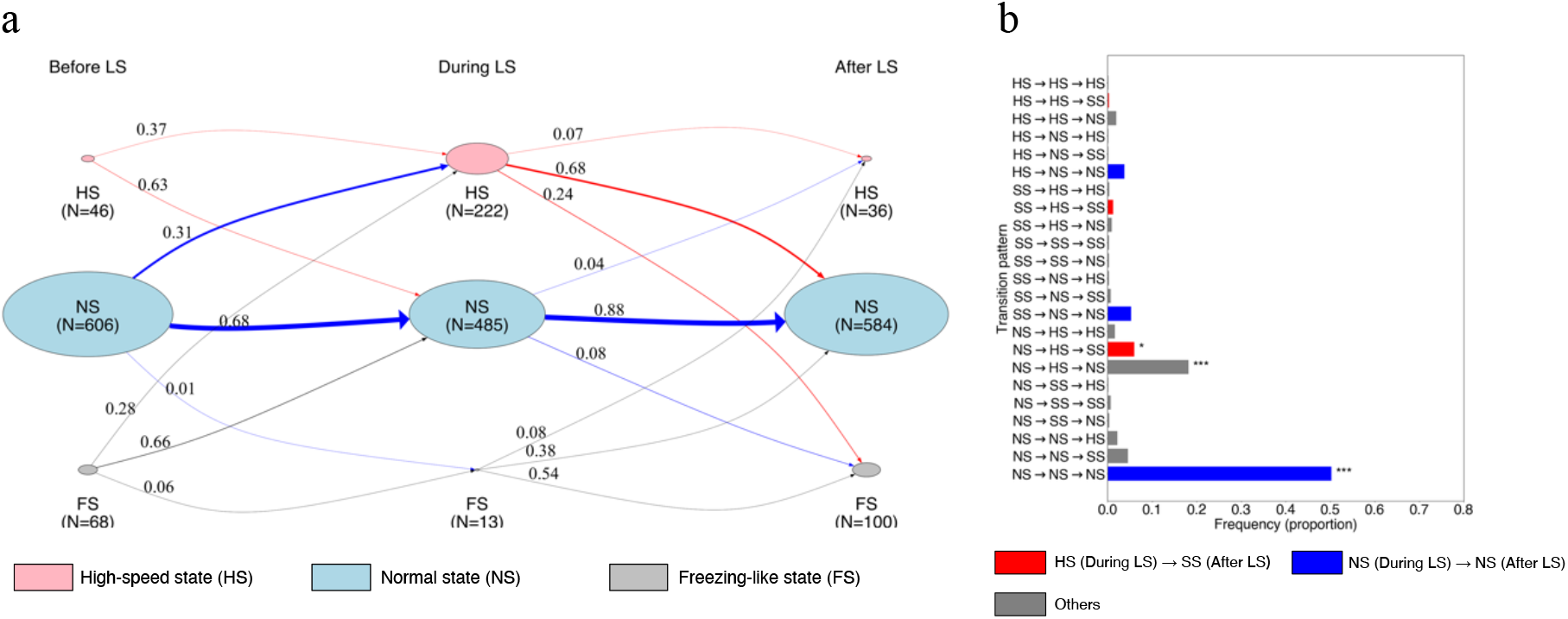
State transition diagram of individual-level behaviour (non-cohabiting group) a) For individual-level behaviours classified as FS: freezing-like, NS: normal swimming, or HS: high-speed swimming, the state transition probabilities were shown from before to during LS and from during to after LS using a Markov chain. Details are as described in Figure 1c. b) Bar graphs showing transition patterns and their frequencies from before LS to during and after LS for FS, NS, and HS. Statistical significance is denoted as follows: ^***^q < 0.001, ^**^q < 0.01, ^*^q < 0.05, and no notation for q > 0.05. Details are as described in Figure 1d.

In Cluster 0 (Figure 3a), the probability of transitioning from the normal state to the high-speed state from before LS to during LS was 72.5%, while the transition probability from the freezing-like state to the high-speed state was 76%. Individuals that exhibited high-speed swimming during LS had an 89% probability of subsequently transitioning to the freezing-like state. In addition, even individuals that remained in the normal state during LS had an 80.4% probability of transitioning to the freezing-like state afterwards. These results suggest that in Cluster 0, both responsive (high-speed swimming) and unresponsive (normal swimming) individuals tended to synchronise into a freezing-like state after LS. The most frequent pattern was NS→HS→FS (red; p < 0.001), and the second was FS→HS→FS (red; q < 0.001), both involving a transition to high-speed swimming during LS followed by freezing-like behaviour (Figure 3d). These findings suggest that Cluster 0 corresponds to groups in which all individuals synchronise and exhibit freezing-like behaviour after LS.

In Cluster 1 (Figure 3b), the probability of maintaining the normal state from before LS to during LS was 86.2%, and individuals that remained unresponsive during LS continued normal swimming after LS with a probability of 94.2%. The dominant patterns were those where individuals maintained the normal state throughout (NS→NS→NS and FS→NS→NS, q < 0.001; Figure 3e, blue). These results suggest that Cluster 1 corresponds to groups in which all individuals synchronise and maintain normal swimming after LS. In the Cluster 2 (Figure 3c and f), NS→NS→NS remained the most frequent pattern (q < 0.001). However, various other transitions were also observed, including NS→HS→NS (q < 0.001; grey), NS→HS→FS (q < 0.001; red), and NS→NS→FS (q < 0.001; grey). This diversity of transitions indicates that Cluster 2 represents a heterogeneous group response profil**e**, reflecting a mixture of multiple individual-level response types rather than a single dominant pattern.

### Synchronisation of freezing-like and non-freezing-like states across the group profiles

Individual-level state transition analysis revealed that in Cluster 0, individuals frequently responded to the looming stimulus (LS) and then entered a freezing-like state, whereas in Cluster 1, transitions in which individuals did not respond to LS and continued swimming were predominant. We next examined whether all individuals in Cluster 0 synchronised to exhibit freezing-like behaviour, and whether all individuals in Cluster 1 synchronised to continue normal swimming. To this end, we counted the number of individuals in the freezing-like state after LS for each group and compared these counts across clusters. A significant difference was observed in the number of individuals exhibiting freezing-like behaviour (Figure 2h, p < 0.001). In Cluster 0, the most frequent outcome was all six individuals showing freezing-like behaviour, while in Cluster 1, the most frequent outcome was none. In Cluster 2, the number of freezing-like individuals ranged mostly from zero to three, yielding a distribution distinct from both Clusters 0 and 1. These results indicate that in Cluster 0, individuals tended to synchronise to the freezing-like state, whereas in Cluster 1 they synchronous to the non-freezing-like state (continued swimming). In contrast, Cluster 2 showed no clear synchronisation, with only a subset of individuals exhibiting freezing-like behaviour after LS.

### Individual-level differences in behavioural transition patterns between cohabited and non-cohabited groups

We determined the behavioural patterns of individuals in the non-cohabited group (Figure S7-8), constructed a state transition diagram (Figure 3a), and classified individual-level transition patterns into 27 categories, comparing their frequencies of occurrence (Figure 3b). As in the cohabited groups, the most frequent pattern was the non-reactive type (NS→NS→NS, blue; q < 0.001). The pattern (NS→HS→NS, grey) in which fish transitioned to a high-speed state during LS and returned to a normal state afterwards also appeared at a significantly high frequency (q < 0.001). In addition, the pattern in which fish transitioned to a high-speed state during LS and then entered a freezing-like state after LS (NS→HS→FS, red) occurred significantly more often (q < 0.05). However, the patterns in which individuals entered a freezing-like state after LS (NS→HS→FS, red; NS→NS→FS, grey), which were significantly enriched in the cohabited groups, did not reach significance in the non-cohabited group. These findings suggest that individual-level transition patterns differed between the two conditions.

### Absence of group-level synchronous freezing-like state in the non-cohabited group

To test whether individuals exhibited synchronous responses after LS, we counted the number of individuals in a freezing-like state per trial for each group and compared the observed data with control data generated by virtual shuffling, as in the cohabited groups (Figure S9). In the observed data, the number of freezing-like individuals peaked at zero, and there was a significant difference in both the number and frequency of freezing-like individuals between the observed and virtual data (χ^2^ (6) = 22.1, p < 0.01) (Figure S9). These results indicate that the peak at zero was not coincidental, but rather that all individuals within the non-cohabited group tended to exhibit synchronous non-freezing behaviour, continuing to swim after LS.

### Differences in synchronous freezing-like states between cohabited and non-cohabited group

To examine whether the occurrence of freezing-like states at the group level after LS differed depending on cohabitation, the number of individuals exhibiting freezing-like states per trial was counted for each group. The aggregated group-level data were then compared between the cohabited and non-cohabited groups using a generalised linear mixed model (GLMM) (Figure S10). The results showed that, following LS, the cohabited groups tended to have a higher number of individuals exhibiting freezing-like states (Figure S10, β = 2.02, p < 0.05) and displayed a bimodal distribution. In contrast, the non-cohabited group showed fewer freezing-like individuals and exhibited a unimodal distribution. This indicates that, unlike the cohabited groups, the non-cohabited group lacked the peak where all six individuals exhibited freezing-like states after LS.

### Disappearance of the collective freezing in the non-cohabited group

To clarify similarities and differences in collective behavioural patterns between cohabited and non-cohabited groups, we integrated and analysed data from both conditions. Specifically, we performed dimensionality reduction and clustering based on 27 individual-level behavioural transition patterns. Principal component analysis (PCA) revealed that 17 dimensions were required to explain 95% of the variance, which was therefore set as the optimal number (Figure S11). The silhouette coefficient indicated that the optimal number of clusters was three (Figure S12), and all groups were classified accordingly (Figure 4a). Groups in the cohabitation condition were distributed across all three clusters, whereas the non-cohabited groups were absent from Cluster 0, indicating a clear bias (Figure 4b). We next verified that in Cluster 0, all individuals tended to exhibit synchronous freezing-like behaviour, while in Cluster 1 they tended to exhibit synchronous non-freezing behaviour (continued swimming). In Cluster 2, synchrony was absent, with only a subset of individuals showing freezing-like behaviour after LS. Importantly, the classification showed that the non-cohabited groups were not represented in Cluster 0, showing that the ‘overall freezing-like state’ cluster was absent from their collective behavioural profiles (Figure 4c).

**Figure 4.**
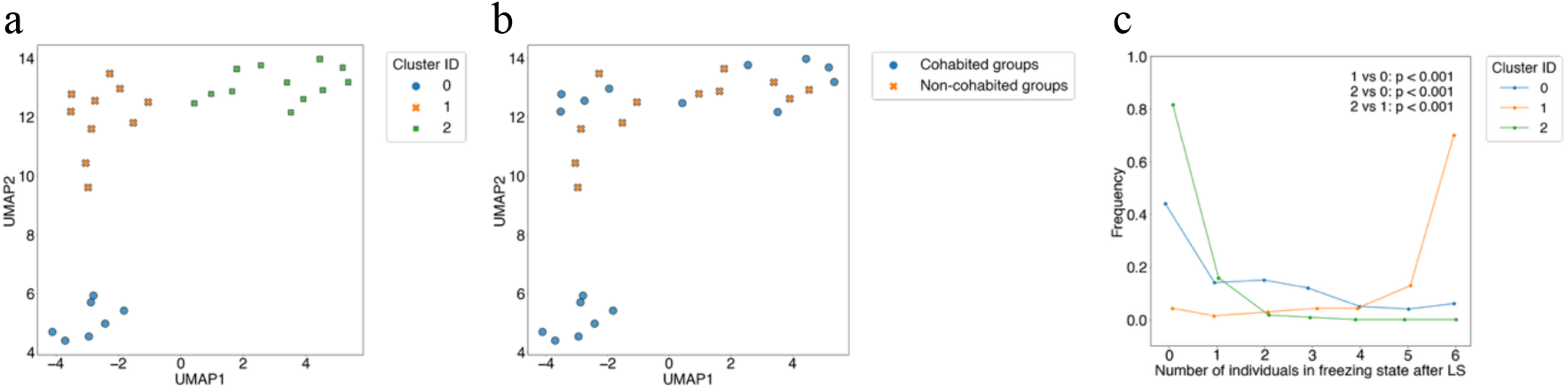
Visualisation of dimensionality reduction using PCA and UMAP, and comparison of freezing-like states across clusters. (a–b) The X-axis represents the first UMAP component and the Y-axis the second. Each point corresponds to the group centroid. (a) Colours indicate cluster IDs obtained by spectral clustering. (b) Colours indicate cohabitation status: cohabitation groups (blue) and non-cohabitation groups (orange). (c) Distribution of freezing-like states (FS) after LS exposure in the integrated dataset combining cohabitation and non-cohabitation groups. Details are as described in Figure 2h.

## Discussion

In this study, we established a quantitative behavioural assay to analyse collective decision-making in medaka (O*ryzias latipes*) in response to a looming stimulus (LS). In this system, small groups of medaka were presented with an LS that mimicked an approaching predator, and their collective behavioural choices were examined. Two dichotomous collective response patterns consistently emerged at the group level: an ‘all-freezing’ state and an ‘all-swimming’ state. Furthermore, the distribution of the number of freezing-like individuals per trial was bimodal, with clear peaks at either zero or six individuals. These results demonstrate the presence of a dichotomous collective behavioural choice in medaka and validate as a robust tool for investigating collective decision-making under controlled laboratory conditions. Moreover, this assay will provide a platform for elucidating the genetic and neural bases of collective decision-making in vertebrates.

Previous studies of collective decision-making under laboratory conditions have primarily used small fish species, such as sticklebacks and golden shiners^2,4,6^, often focusing on wild populations in ecological contexts. By contrast, our study employed medaka, a well-established genetic model organism, thereby enabling experimental systems in which genetic and environmental factors can be controlled. This approach enables the establishment of highly reproducible behavioural assays and allows for long-term monitoring of behavioural development, from the individual to the group level.

Most previous studies of collective decision-making have focused on gradual responses to predators^6,7^. In contrast, our findings revealed a novel phenomenon: rapid collective responses to sudden visual threats. In coral reef fishes, escape responses of individuals can be predicted from the expansion rate of looming stimuli or the behaviour of neighbours^24^. However, how these individual responses converge into synchronous group-level behaviour remains unclear. Our results demonstrate that under time-constrained predatory threat, rapid collective decision-making can emerge, thereby complementing existing models of gradual escape behaviour.

Our findings further suggest that a certain period of cohabitation is required for collective behavioural choices in response to the LS. In particular, in cohabited groups, many individuals transitioned from high-speed swimming during LS to freezing afterwards, and entire groups tended to enter the freezing-like state. Such consistent behavioural synchrony was mainly observed in groups that had undergone sufficient cohabitation, suggesting that social familiarity may contribute to coordinated collective decisions. In our experiments, cohabitation increased the proportion of individuals that exhibited a freeze after escape-like high-speed swimming, indicating a change in behavioural regularity at the individual level. However, these individual-level changes alone cannot fully explain the emergence of dichotomous collective outcomes (all-freezing versus all-swimming). It remains unclear whether cohabitation (1) enhanced each individual’s social sensitivity, making them more likely to be influenced by others, or (2) homogenised behavioural traits within groups. Distinguishing between these two possibilities was beyond the scope of this study. Future approaches incorporating longitudinal tracking of individually identified fish and quantitative measures of behavioural synchrony will be necessary to address this question.

If explanation (1) is correct, repeated interactions during cohabitation may allow individuals to recognise and predict the behaviour of conspecifics, thereby strengthening group-level properties such as polarisation and alignment. Previous studies have reported various effects of social familiarity in fish. For example, in female guppies, 12 days of cohabitation led to preferential associations with familiar conspecifics^31^. Social familiarity has also been shown to promote group cohesion and alignment in guppies^14^ and to enhance information transfer under social threat in damselfish^17^10. In addition, cohabitation may also induce social fear contagion. In zebrafish, individuals are known to switch from high-speed swimming to freezing when exposed to alarm cues from conspecific skin extracts^13,32^, suggesting that this behavioural pattern may be widespread among fishes. Moreover, zebrafish exhibit similar freezing responses when observing familiar conspecifics or groups displaying fear responses^13,33^.

On the other hand, explanation (2), that cohabitation homogenises behavioural traits within groups, cannot be excluded. Previous studies have shown that bold individuals tend to maintain stable behavioural traits, whereas shy individuals are more plastic and influenced by social context. For instance, in guppies, bold individuals rely on their own information and explore independently, whereas shy individuals adjust their behaviour according to social information^34^. In sticklebacks, bold individuals also show stable exploratory behaviour, while shy individuals display behavioural plasticity and can change over time^35^. However, to our knowledge, no studies have directly demonstrated long-term homogenisation of behavioural traits caused by cohabitation in any animal species. Thus, we consider explanation (1) to be the more plausible mechanism underlying our findings.

In summary, our results indicate that cohabitation promotes the dichotomisation of collective behavioural choices in medaka, and that factors such as social familiarity or changes in social sensitivity may contribute to this process. Although further investigation will be required to directly verify these mechanisms, our study provides a foundation for exploring how social experience shapes collective decision-making under time-constrained predatory threats in vertebrates.

## Acknowledgements

We thank Tetsuro Takeuchi for sharing the fading strain. We thank Drs. Masahiro Daimon and Masayuki Koganezawa for their advice on the development of the behavioural assay. We thank Profs. Ken-Ichiro Tsutsui, Hiromu Tanimoto, Jamie M. Kass and Dr. Towako Hiraki-Kajiyama for comments on the manuscript. This work was supported by the National Institute for Basic Biology Priority Collaborative Research Project 10-104 (to H.T.), 19-347 (to H.T.), and 21-335 (to H.T.); a grant for Joint Research (#01111904) by the National Institutes of Natural Sciences (to H.T.); Japan Society for the Promotion of Science (JSPS) KAKENHI Grants 21H04773 (to H.T.), 20H04925 (to H.T.), 18H02479 (to H.T.), 22H05483 (to H.T.), 23K27205 (to H.T.), 24H01216 (to H.T.) and 24K21957 (to H.T.). Takeda Science Foundation (to H.T.), and the natural science grant of the Mitsubishi Foundation (to H.T.); Japan Society and Technology Agency (JST) SPRING, Grant Number JPMJSP2114(to R.N.).

## Funding

This work was supported by the National Institute for Basic Biology Priority Collaborative Research Project 10-104 (to H.T.), 19-347 (to H.T.), and 21-335 (to H.T.); a grant for Joint Research (#01111904) by the National Institutes of Natural Sciences (to H.T.); Japan Society for the Promotion of Science (JSPS) KAKENHI Grants 21H04773 (to H.T.), 20H04925 (to H.T.), 18H02479 (to H.T.), 22H05483 (to H.T.), 23K27205 (to H.T.), 24H01216 (to H.T.) and 24K21957 (to H.T.). Takeda Science Foundation (to H.T.), and the natural science grant of the Mitsubishi Foundation (to H.T.); Japan Society and Technology Agency (JST) SPRING, Grant Number JPMJSP2114(to R.N.).

## Author Contributions

R.N. and H.T. designed experiments. R.N. conducted experiments, wrote code and analysed data. R.N. and H.T. co-wrote and edited the paper and supervised the project.

## Competing interests

The authors declare no competing interests.

## Data availability

All data generated or analysed during this study are available from the corresponding author on reasonable request.

## Notes

### Competing Interest Statement

The authors have declared no competing interest.

### Summary of Updates

This revised manuscript represents a substantial update that supersedes the previous posting. The scope has been reframed to test directly whether cohabitation shapes instantaneous collective decision-making under looming threat. To this end, we performed new experiments that included non-cohabited groups assembled immediately before testing, allowing a direct contrast with cohabited groups. Methods and analyses were extensively restructured. The tracking and preprocessing pipeline was standardised; behavioural state definitions across pre, during, and post stimulus intervals were clarified; and state transition matrices were computed. At the group level, response profiles were defined using dimensionality reduction (PCA followed by UMAP) and spectral clustering. Group outcomes were evaluated with generalised linear mixed models (GLMMs) with appropriate control for multiple comparisons. The key results were updated. Whereas the previous version reported a dichotomy between all-freezing and all-swimming group outcomes, this revision newly identify a third partial-freezing profile and further show that the all-freezing profile is absent in non-cohabited groups. All main figures and supplementary files were replaced to reflect these new analyses and datasets. The Abstract, Introduction, Results, Methods, and Discussion were comprehensively rewritten for clarity and completeness. The title was updated to reflect the new emphasis, and author affiliations, acknowledgements, funding information, and the data availability statement were revised accordingly. Overall, this revision replaces most analyses and figures, provides new experimental evidence and establishes a strengthened quantitative framework.

