## Supplemental Figure for "Cohabitation Shapes Collective Decision-Making in Medaka Fish"

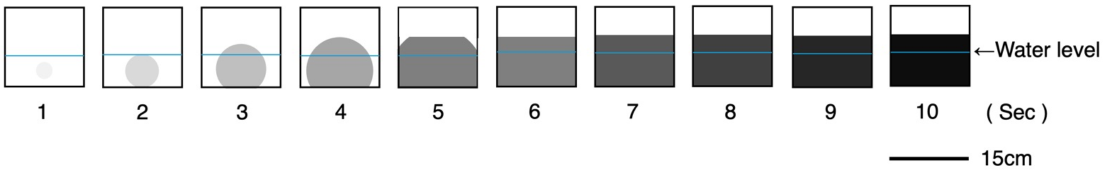


Figure S1 Schematic representation of the looming stimulus at 1 frame per second

Each rectangle represents the looming stimulus (LS) presented on the side of the tank via the display at 1-second intervals. The expansion of the circle is completed in 5.5 seconds, followed by a gradual darkening over the subsequent 10 seconds. The blue horizontal line and the numbers below each rectangle indicate the water level and the elapsed time of the LS, respectively. The water level was adjusted to approximately 6 cm.

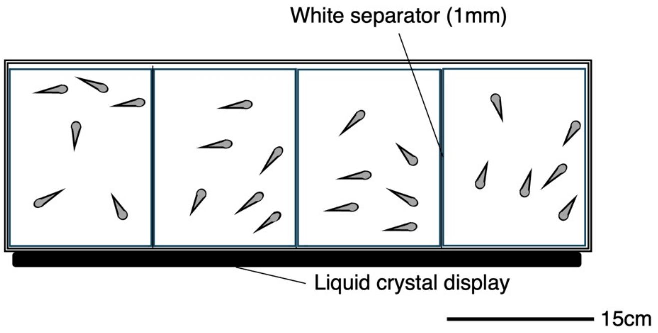


Figure S2 Schematic representation of the behavioural experimental apparatus viewed above

To prevent visual interference, partition boards were placed between each tank. Six fish were placed in each tank, and the apparatus allowed for simultaneous testing of up to four tanks. The looming stimulus (LS) was presented simultaneously to the four tanks using a liquid crystal display. The entire apparatus was enclosed by white panels to block external stimuli.

a

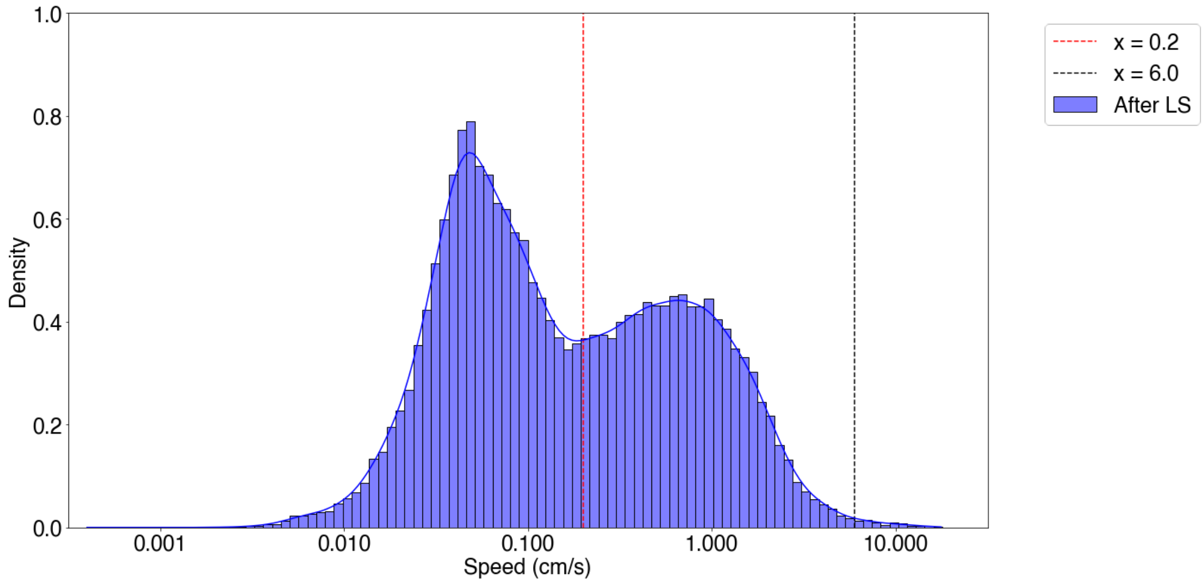


b

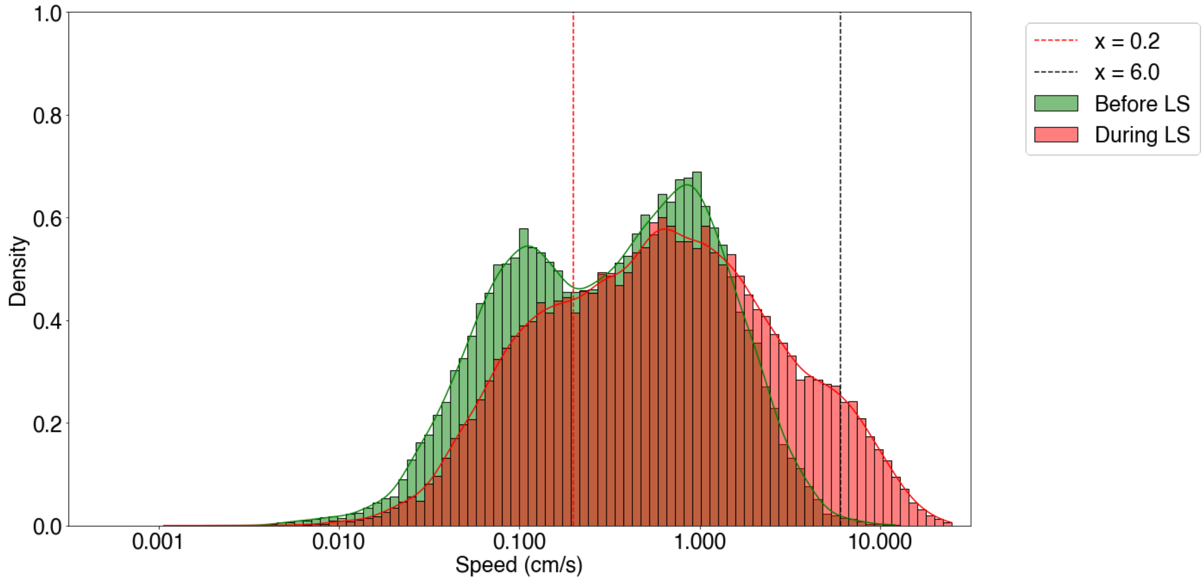


Figure S3 Distribution of speed per frame (Cohabited group)

The x-axis represents speed, and the y-axis represents density. The figure shows histograms with a bin width of 0.05 along with curves obtained by kernel density estimation (KDE). Vertical dashed lines denote speeds of 0.2 cm/s (red) and 6.0 cm/s (black). (a) Distribution of speed after LS (blue). (b) Distribution of speed before (green) and during (red) LS.

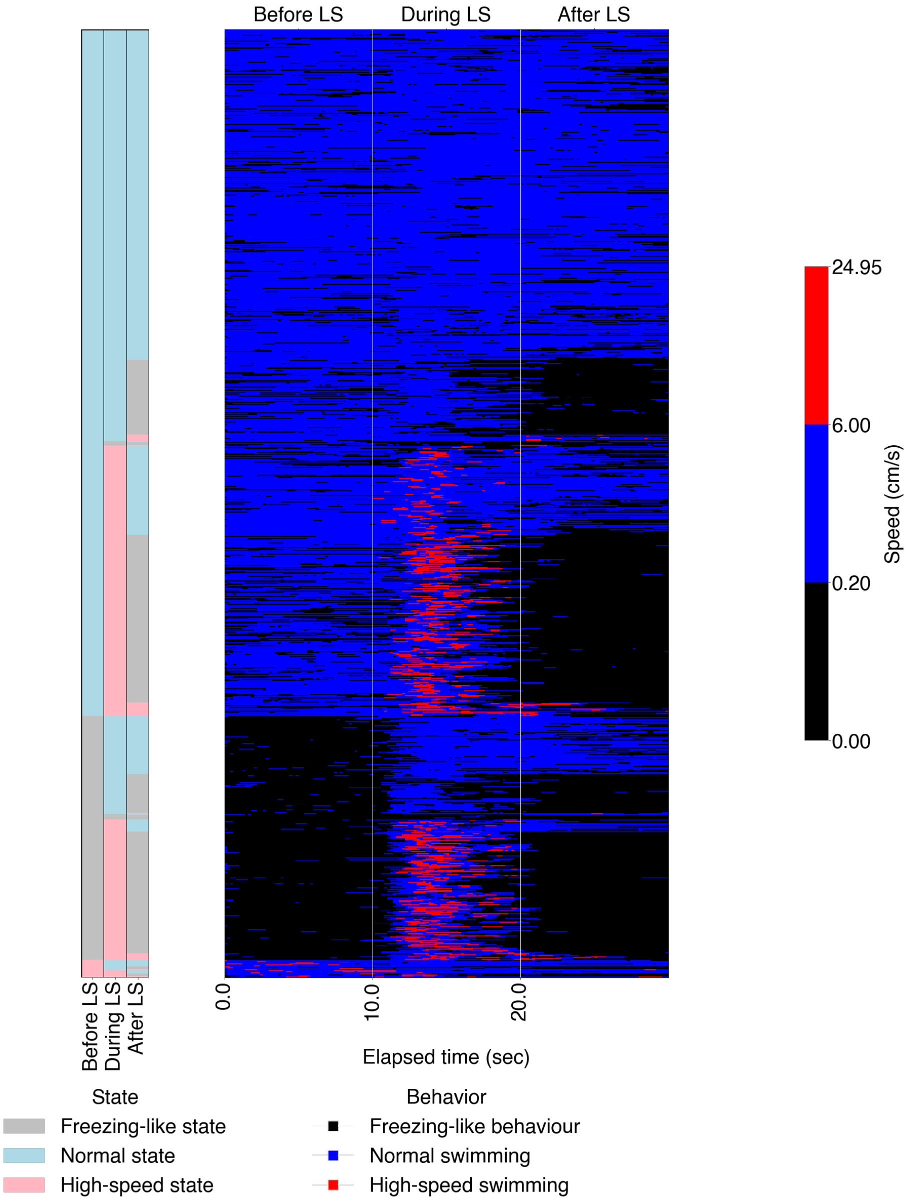


### Figure S4 Heatmaps visualizing behavioural classification and behavioural patterns (Cohabited group)

In the heatmaps, the x-axis represents elapsed time, and the y-axis represents individual data for each trial. The colour bar on the right indicates the classification of three behaviours for each frame. The colour bar on the left indicates the behavioural classification for the three periods: before, during, and after the stimulus. Grey, light blue, and pink indicate a freezing-like state, a normal state, and a high-speed state, respectively. (a) In the heatmap cells, black indicates freezing-like behaviour, blue indicates normal swimming, and red indicates high-speed swimming. Freezing-like behaviour was defined as frames with speed below 0.2 cm/s; normal swimming as frames with speed between 0.2 cm/s and 6 cm/s; and fast swimming as frames with speeds of 6 cm/s or higher.

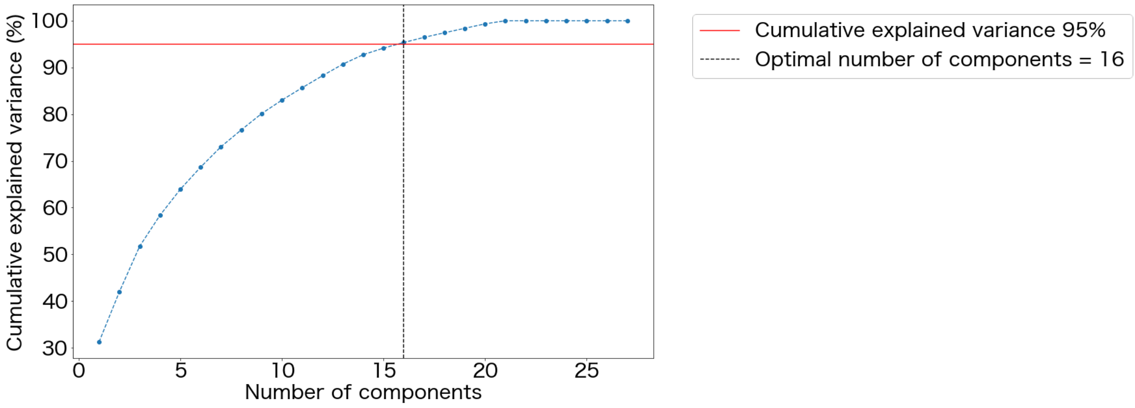


### Figure S5. Cumulative contribution rate and number of principal components in the cohabitation group

This figure shows the cumulative contribution rate obtained when reducing the dimensions of a matrix summarising 27 individual behavioural patterns per group. The X-axis represents the number of principal components, and the Y-axis represents the cumulative contribution rate. The red horizontal line indicates a cumulative contribution rate of 95%. Based on this criterion, the optimal number of principal components was 16.

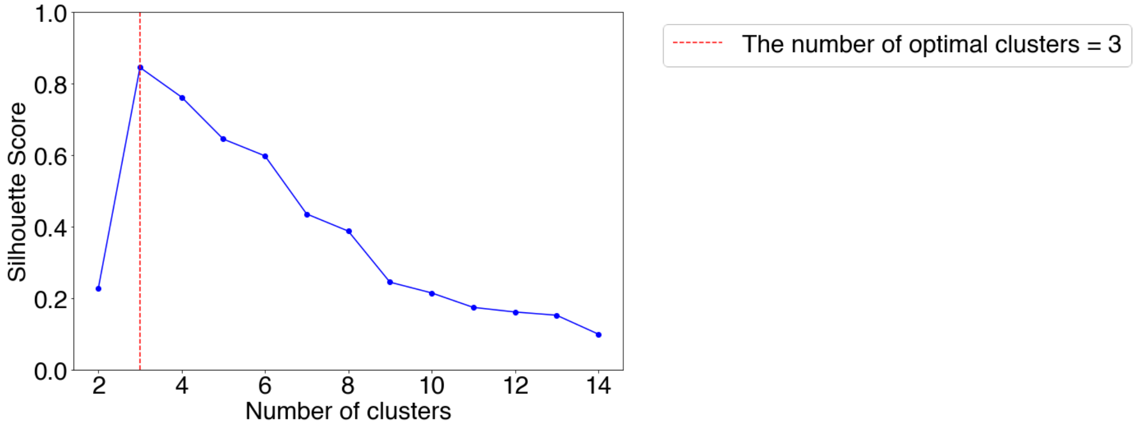


### Figure S6. Silhouette coefficient and number of clusters in the cohabitation groups

The X-axis represents the number of clusters, and the Y-axis represents the silhouette coefficient. A value close to 1 indicates strong cohesion within clusters and clear separation between clusters. Spectral clustering yielded the highest silhouette coefficient (0.845) when the number of clusters was set to three, indicating that the optimal number of clusters was three.

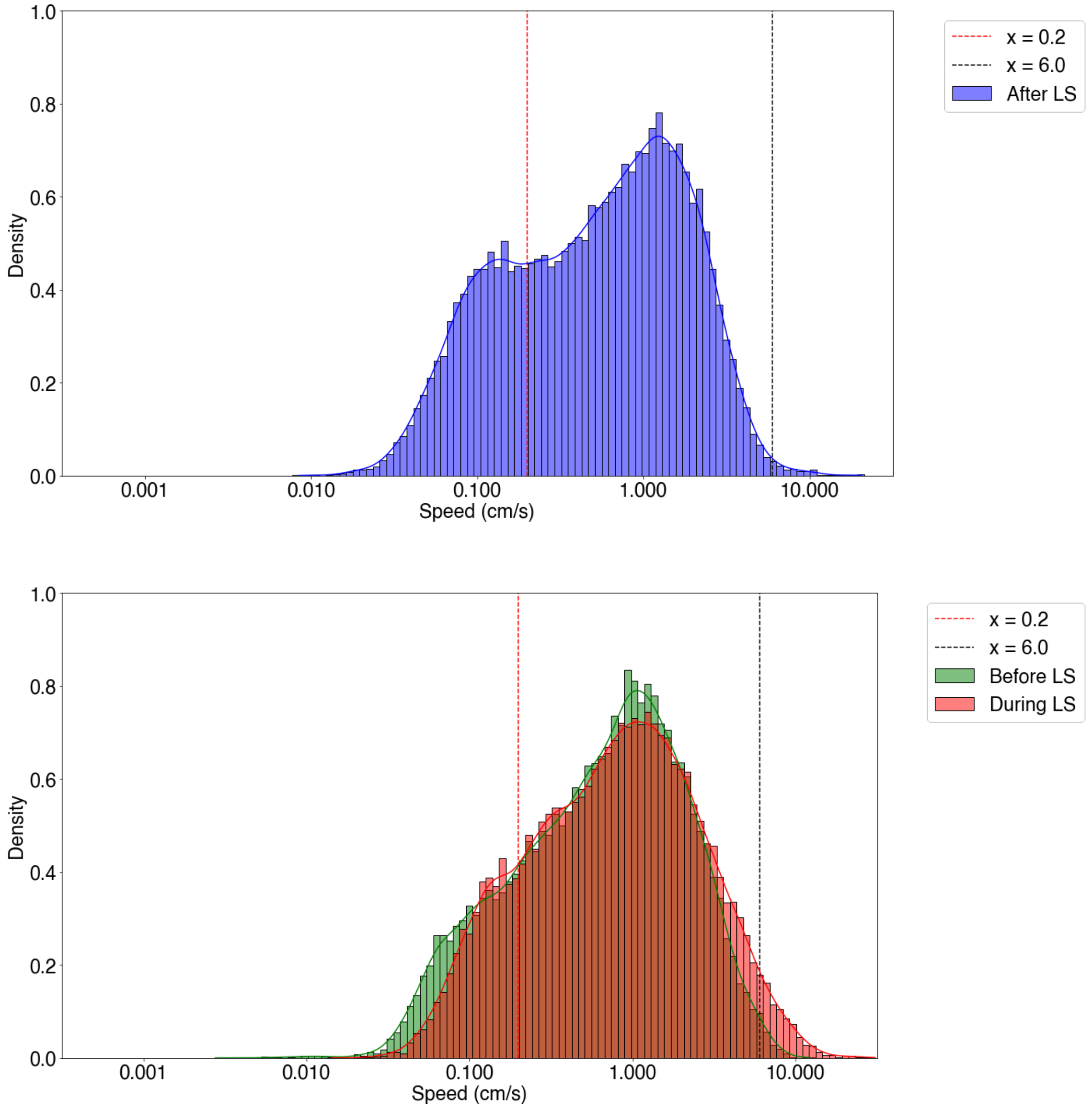


Figure S7. Distribution of frame-by-frame swimming speed in the non-cohabitation group.

Details are as described in Figure S3. (a) Speed distribution after exposure to the LS (blue). (b) Speed distributions before (green) and during (red) the LS.

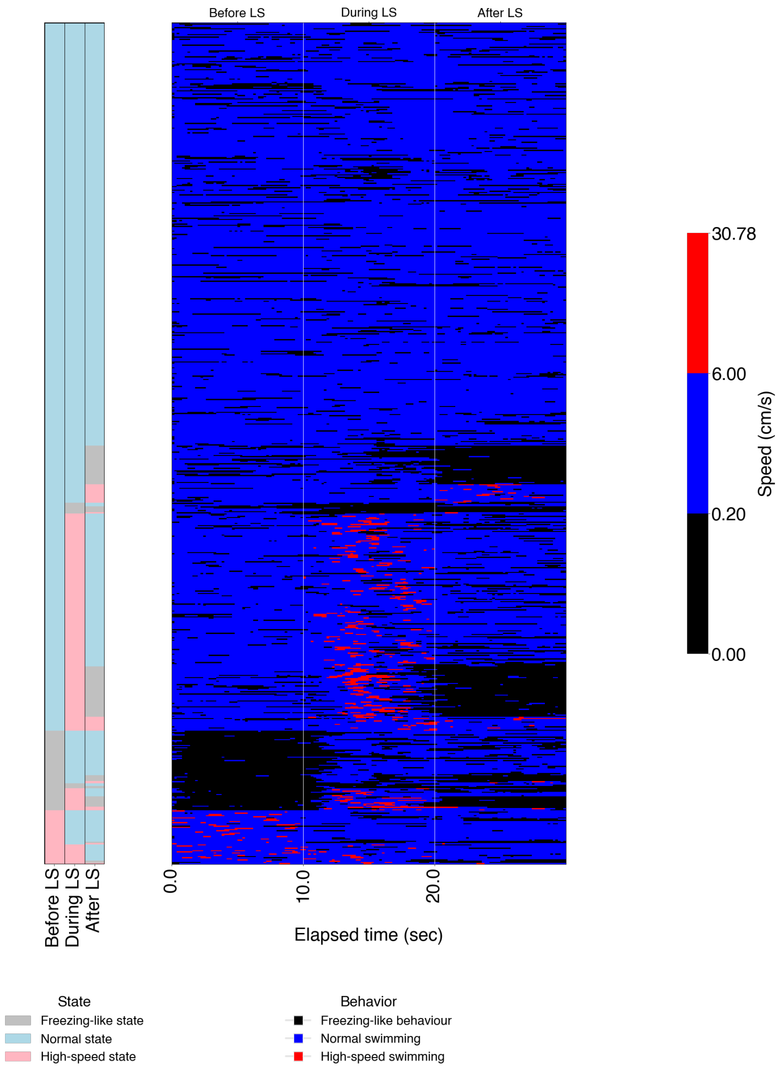


### Figure S8. Heatmaps visualising behavioural classification and patterns in the non-cohabitation group

The structure of this figure is the same as in Figure S6, but it shows data from the non-cohabitation group.

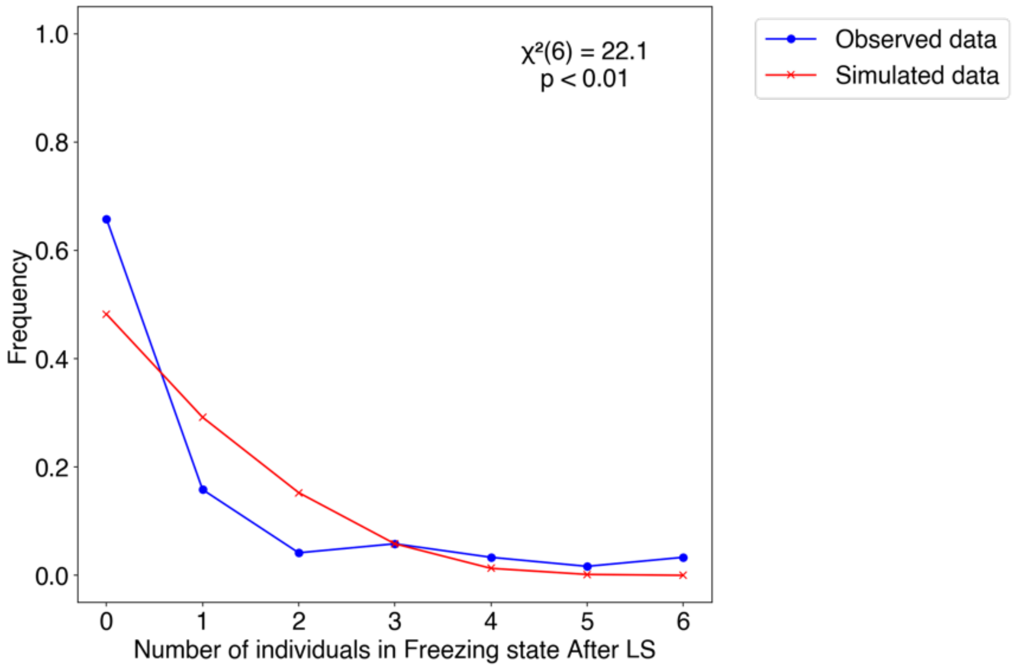


### Figure S9. Distribution of the number of individuals in a freezing-like state after LS (Non-cohabited group)

The X-axis shows the number of individuals in the freezing-like state, and the Y-axis shows the relative frequency across all trials. The blue line represents the observed data (12 groups), and the red line represents the simulated data (12 groups × 1000 trials, seeds 1–1000). A chi-square test was conducted, and the results are shown in the graph ($\chi^2$ (6) = 22.1, $p < 0.01$). The degrees of freedom, test statistic, and p-value are indicated.

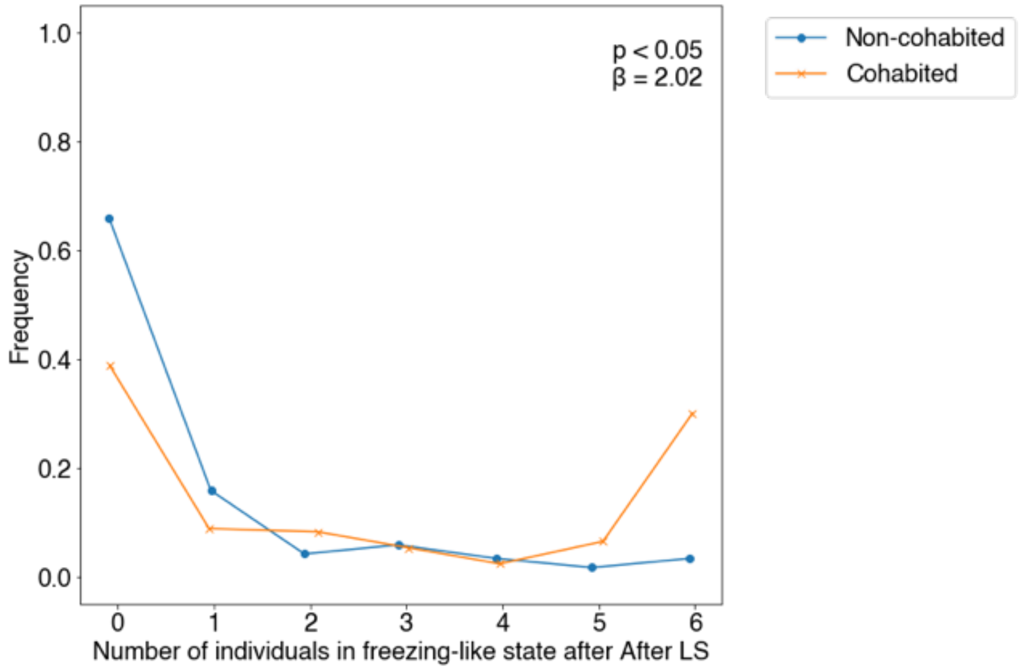


Figure S10 Effect of cohabitation on the distribution of individuals in freezing-like state after LS

The blue line corresponds to the non-cohabited group, and the orange line corresponds to the cohabited group. Results of the GLMM are indicated in the figure legend. β denotes the regression coefficient. A significant positive coefficient indicates that the number of individuals in the freezing-like state was significantly higher in the cohabitation group.

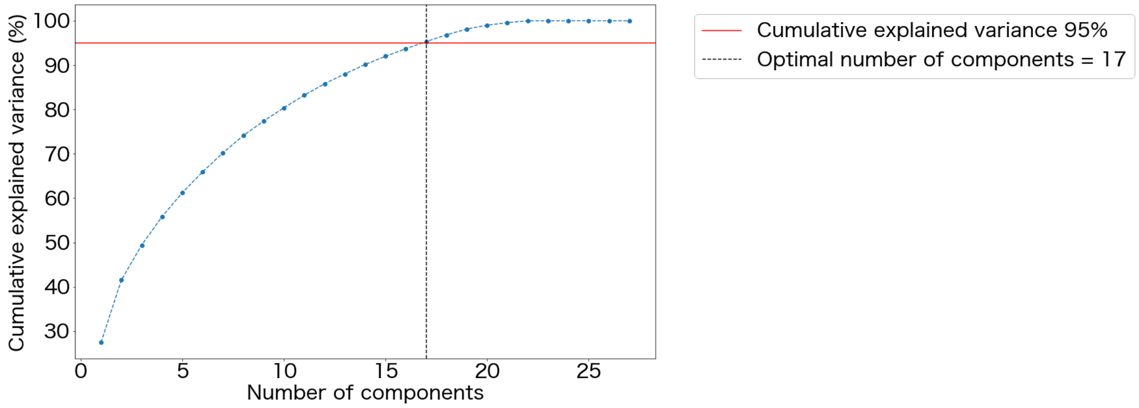


Figure S11. Cumulative contribution rate and number of principal components in the cohabitation group

Details are as described in Figure S5. Based on this criterion, the optimal number of principal components was 17.

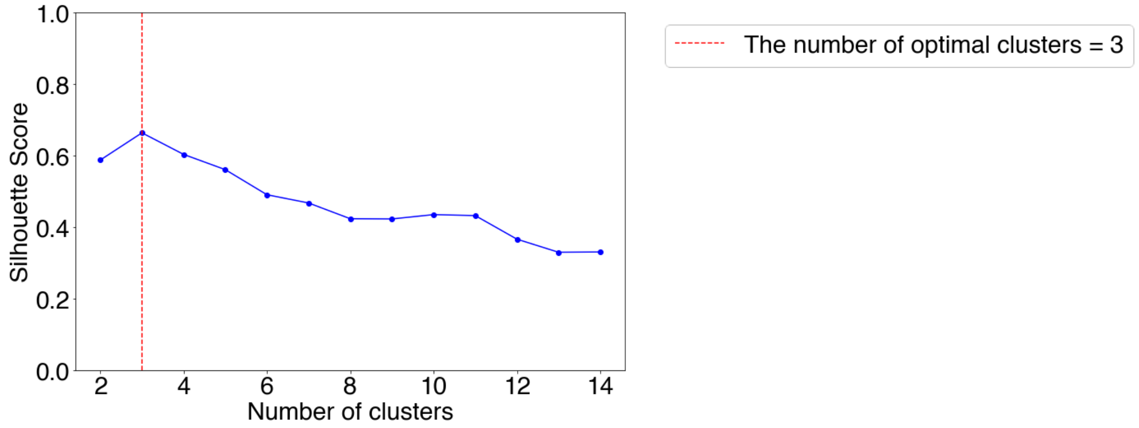


Figure S12. Silhouette coefficient and number of clusters in the cohabitation group

Details are as described in Figure S6. Spectral clustering yielded the highest silhouette coefficient (0.663) when the number of clusters was set to three, indicating that the optimal number of clusters was three.
